# An improved molecular tool for screening bacterial colonies using GFP expression enhanced by a *Dictyostelium* sequence

**DOI:** 10.1101/821157

**Authors:** Tomo Kondo, Shigehiko Yumura

**Affiliations:** Graduate School of Sciences and Technology for Innovation, Yamaguchi University, 753-8512 Yamaguchi, Japan

## Abstract

During molecular cloning, screening of bacterial transformants is a time-consuming and labor-intensive process. However, tractable tools that can be applied to various vectors for visual confirmation of desired colonies are limited. Recently, we reported that TED (translational enhancement by a *Dictyostelium* gene sequence) boosted protein expression even without an expression inducer in *Escherichia coli*. Here, we demonstrate a generally applicable molecular tool using the expression of green fluorescent protein (GFP) enhanced by TED. By inserting a module related to TED into the cloning site in advance, we effectively screened *E. coli* colonies harboring the desired plasmid functions in a prokaryote (*Magnetospirillum gryphiswaldense*) or eukaryote (*Dictyostelium discoideum*). Thus, our system represents a user-friendly technique for cloning.

## Introduction

Molecular cloning is a technique used to create DNA constructs for studies in various biological fields, including molecular cell biology, biotechnology, and synthetic biology. Because of extensive studies using *Escherichia coli*, there are many versatile methods available for molecular cloning (1). However, screening methods for bacterial transformants that contain the desired plasmid are limited.

One of the popular color-based screening methods is blue-white screening using the pUC vector series (2, 3). The vector contains the α-peptide gene (4) in the multi-cloning site (MCS). After its translation, followed by complementation of the β-galactosidase, blue pigments are produced through hydrolysis of the colorless lactose analogue 5-bromo-4-chloro-3-indolyl-β-D-galactoside (X-gal). When the α-peptide gene is disrupted by insertion of a DNA fragment into the MCS, the colony appears white. Because α-peptide expression is controlled by the *lac* operon, it can be upregulated by the addition of isopropyl β-D-thiogalactopyranoside (IPTG) (5). Although this method is simple, various factors cause false-positive results, including the presence of an intact α-peptide gene without corresponding product synthesis (1). Therefore, these false-positive results often result in unnecessary costs and labor because superfluous colonies are handled in the next procedure.

Alternatively, fluorescence-based screening has been developed for colony screening (6–11) and is based on the functionality of the green fluorescent protein (GFP), even in *E. coli* (12). A nonfluorescent colored protein (i.e., chromoprotein) has also been used (13). Using these visual markers, the screening enables efficient selection of colonies containing the desired vector inserts. However, these protein expressions often require an expression inducer such as IPTG, and its general feasibility for a variety of vectors has not yet been reported.

Here, we present a tractable tool for bacterial colony screening by GFP fluorescence based on a system called TED that enhances protein expression using a gene sequence of the social amoeba *Dictyostelium discoideum* in *E. coli* (14). In our previous work, insertion of a *Dictyostelium* gene sequence (e.g., *mlcR*) upstream of the GFP gene can spontaneously induce visible levels of GFP expression without the addition of IPTG. We applied this method to generate shuttle vectors for expression of heterologous proteins in magnetotactic bacteria or *D. discoideum*. We propose to incorporate this tool into a variety of vectors to raise the efficiency of colony screening.

## Materials and methods

### E. coli culture and transformation

*E. coli* K12 strain HST08 Premium (Takara Bio, Shiga, Japan) was used for cloning. Lysogeny broth (LB) medium was obtained from Nippon Genetics (Tokyo, Japan) and supplemented with 100 µg/mL ampicillin. After chemical transformation (15, 16), cells were selectively grown on LB agar prepared by solidifying the LB media with 1.5% agar supplemented with 100 µg/mL ampicillin.

### Magnetotactic bacteria culture and transformation

#### Magnetospirillum gryphiswaldense

MSR-1 was obtained from DSMZ (Braunschweig, Germany). Cells were cultured at 28ºC in DS medium containing 2.38 g HEPES (Dojindo Laboratories, Kumamoto, Japan), 3 g sodium pyruvate (Wako Pure Chemical Industries), 0.1 g yeast extract (Oxoid, Basingstoke, Hants, UK), 3 g soybean peptone (Sigma-Aldrich, St Louis, MO, USA), 0.34 g NaNO_3_ (Wako Pure Chemical Industries), 0.1 g KH_2_PO_4_ (Wako Pure Chemical Industries), 0.15 g MgSO_4_·7H_2_O (Wako Pure Chemical Industries), and 3 mL of 10 mM ferric quinate in 1 L of distilled water, which was slightly modified from a previous study (17). The pH of DS medium was adjusted to 7.0 with NaOH. Ferric quinate (10 mM) was prepared by adding 4.5 g of FeCl_3_·6H_2_O (Kanto Chemical, Tokyo, Japan) and 1.9 g of quinic acid (Sigma-Aldrich) to 1 L of distilled water. For agar-plate culture, activated charcoal (1 g/L, Wako Pure Chemical Industries) was added to the above DS medium. After autoclaving, 1.4-dithiothreitol (final concentration of 1 mM) was added aseptically and solidified by adding 1.5% agar. Antibiotics (5 µg/mL ampicillin) were supplemented, as needed.

Transformation was carried out by electroporation. For each pulsing experiment, 15 mL of exponential phase cells grown in 15 mL polypropylene tubes was collected by centrifugation (1200 × *g* for 10 min) at 4ºC. The cells were washed twice using cold electroporation buffer (1 mM HEPES, pH 7.0, 1 mM MgCl_2_, 272 mM sucrose) and resuspended in about 50 μL of the same buffer. The mixture was loaded in a chilled 0.1-cm cuvette with plasmid DNA and subjected to a single pulse using a Gene Pulser Xcell (Bio-Rad Laboratories, Hercules, CA, USA) set at 25 μF capacitance, 200 Ω resistance, and 0.8–1.0 kV voltage. The cells were recovered in 1 mL of fresh DS medium overnight at 28ºC and then plated onto a DS medium agar plate with antibiotics. For approximately one week, visible colonies were observed. After checking the colonies by PCR using EmeraldAmp MAX (Takara Bio), the selected cells were grown in DS medium with antibiotics. Plasmids were purified using NucleoSpin Plasmid (Macherey-Nagel, Düren, Germany) following the manufacturer’s instructions.

### Dictyostelium culture and transformation

*D. discoideum* cells of the Ax2 were grown in HL5 medium (15.4 g glucose, 7.15 g yeast extract, 14.3 g bacteriological peptone, 0.486 g KH_2_PO_4_, 1.28 g Na_2_HPO_4_·12H_2_O, per liter) at 22ºC (18). Additional antibiotics (G418, blasticidin S, streptomycin, and ampicillin) were supplemented at appropriate concentrations, as needed. Transformation was carried out by electroporation as described previously (19) with modifications. Briefly, cells were washed with electroporation buffer (50 mM sucrose, 10 mM Na-K-phosphate buffer, pH 6.1) and collected by centrifugation (536 × *g* for 2 min) at 4ºC. After removing the supernatant, cells were suspended in about 100 µL of the same buffer. The mixture was loaded in a chilled 0.1-cm cuvette with plasmid DNA and subjected to two square pulses at 5 s intervals using the Gene Pulser Xcell (Bio-Rad Laboratories) set at 0.55 kV voltage. The cells were recovered in HL-5 overnight at 22ºC, and the appropriate antibiotics were added for selection.

### Plasmids

Plasmids used in the study were constructed using standard techniques. For seamless cloning, In-fusion cloning (Takara Bio) was used following the manufacturer’s protocols. The pMGT fragment (20), *msp3* promoter (21), *mms13* (22), and HaloTag gene (23) were synthesized by Genewiz (South Plainfield, NJ, USA). The gene that encodes the HaloTag protein was optimized for *Magnetospirillum* codons with permission from Promega Corporation (Madison, WI, USA). The *neo* gene encodes amino-glycoside 3’-phosphotransferase with the promoter, and the terminator and fragment of Ddp1 (24) were cloned from the pBIG vector (GenBank accession number AF270470) (25). The *actin15* promoter and terminator with MCS were synthesized by Genewiz. The genes blasticidin-S deaminase (*bsd*), mEGFP (referred to as simply GFP), and mRuby3 were synthesized with codon optimization for *Dictyostelium* (FASMAC, Kanagawa, Japan; Genewiz).

### Imaging

For excitation of GFP on an agar plate, a custom-made blue LED illuminator was used. For imaging of magnetotactic bacteria, cells were collected by centrifugation (4300 × *g* for 3 min) at 25ºC and then washed twice with distilled water. The cells were incubated with HaloTag TMR Ligand (1 µM at the final concentration) in distilled water at 25ºC for 1 h in the dark. After washing twice, cells were added onto a coverslip (24 × 60 mm) and overlaid with a block of 1.5% agar (8 × 8 × 1 mm) in distilled water. Fixed *Dictyostelium* cells were prepared as described previously (26). The specimens were observed with a microscope (DIAPHOT300 or ECLIPSE Ti, Nikon) equipped with a camera (Orca-ER C4742-80-12AG, Hamamatsu Photonics; or Zyla 4.2, Andor technology, Belfast, Northern Ireland) and mercury lamp (Nikon) using a 100x objective lens. Fluorescence in magnetotactic bacteria was evaluated using a custom written macro in Fiji (27).

## Results and discussion

### Colony screening using TED-induced GFP expression

We developed a GFP expression vector in combination with TED without any adverse effects such as growth rate (14). In this system, *gfp*, which is placed downstream of the 25-bp fragment of the 3’ end of *mlcR* (*mlcR25*) and the SD sequence, is expressed under control of the *lac* promoter, but IPTG is not necessary. Even in the absence of IPTG, fluorescence can be readily distinguished by excitation with a blue LED. The sequence region necessary for GFP expression is hereinafter referred to as the TED module (Fig. 1A).

**Figure 1.**
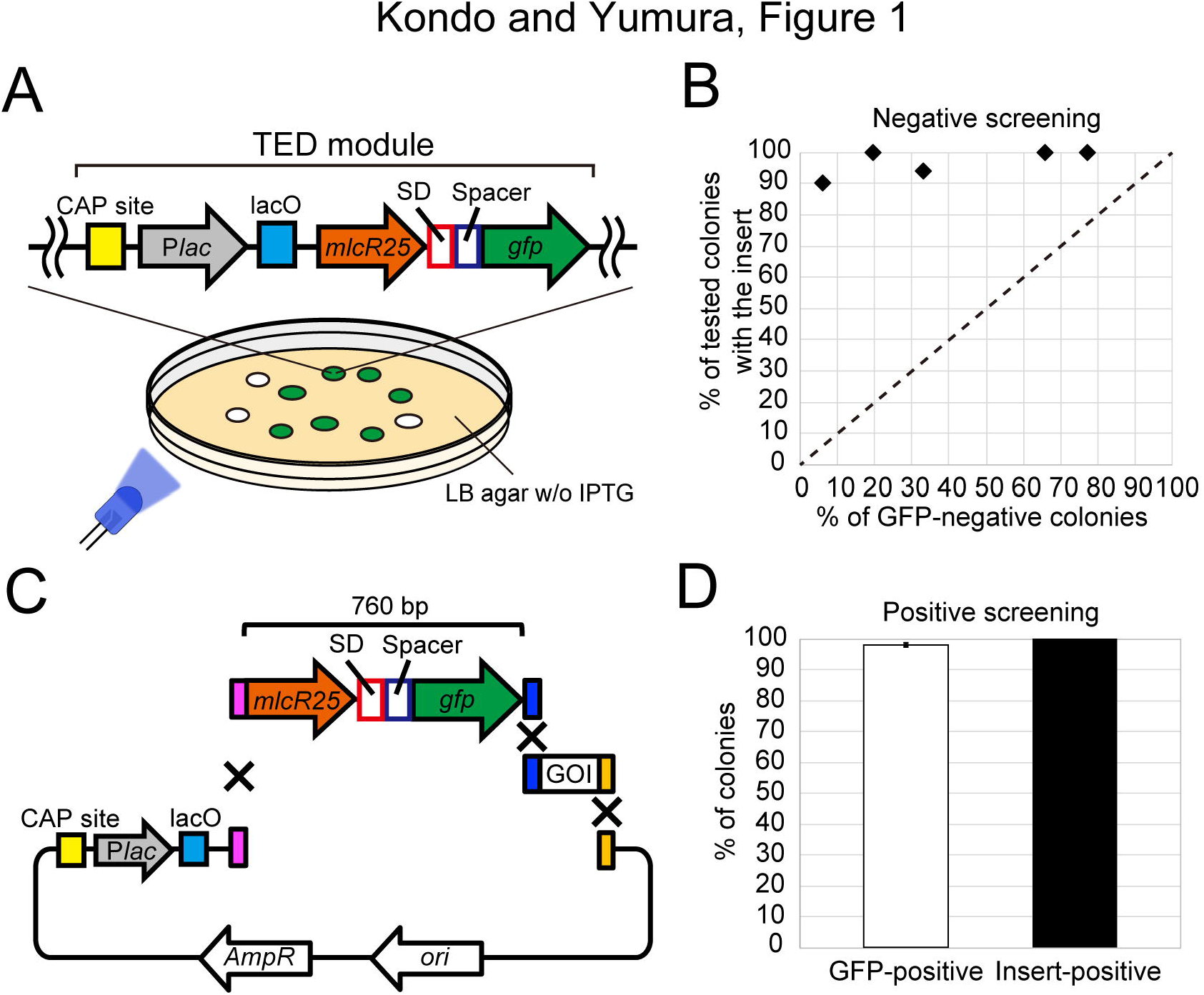
GFP fluorescence-based colony screening. (A) A schematic representation showing colonies with or without GFP expression under blue LED illumination. GFP-expressing colonies carry plasmids containing the TED module, which contains the catabolite activator protein (CAP) binding site, *lac* promoter (P*lac*), *lac* operator (lacO), 25-bp fragment of the 3’ end of *mlcR* (*mlcR25*), SD sequence (SD), spacer, and *gfp*. (B) Success rate of screening colonies with the insert (*tubA*) among GFP-negative colonies. A ligation reaction was carried out with (black rhombuses) or without an insert and used for transformation. The dashed line shows the theoretical success rate when the TED module is not used. In these experiments, less than 3% of colonies were observed as false-positive (GFP-negative without the target insertion). n ≥ 8. (C) A schematic representation depicting the plasmid design for GFP-positive colony selection. *mlcR25* indicates the 25-bp fragment of the 3’ end of *mlcR*. (D) The percentages of GFP-positive colonies (white) and colonies with successful insertion of the target (mRuby3 gene) among GFP-positive colonies (black).

First, we tested a negative screening method to detect successful target insertion by the loss of GFP fluorescence. Part of the TED module (*mlcR25*-SD sequence-spacer-*gfp*) in the pUC19 vector was replaced with an *Xba*I-*Bam*HI fragment of the target sequence encoding *D. discoideum* α-tubulin (1395 bp) as a representative. We obtained plates with various ratios of GFP-negative (i.e., target-inserted) and GFP-positive (i.e., empty vector) colonies by varying the insert concentration used for DNA ligation (Fig. 1B, x-axis). We found that more than 90% of GFP-negative colonies were successfully inserted in any condition (Fig. 1B, y-axis). For negative controls, without addition of the insert with the vector in the ligation mixture, GFP-negative colonies were observed at a rate of ~3%. Under these conditions, even for a ‘worst’ plate with 6% of colonies showing no GFP fluorescence, we could select colonies containing the target sequence with 90% probability. Thus, these results suggest that this method has sufficient sensitivity and high reliability.

We also tested a positive screening method to detect successful target insertions by the appearance of GFP fluorescence. Seamless cloning is a powerful method to generate constructs. However, there are few tools for visually confirming desired colonies on the plate, except for antibiotic selection. Thus, we seamlessly cloned the TED module into the pUC19 vector with the sequence of a gene-of-interest (GOI) (Fig. 1C). In this case, 98 ± 0.6% of colonies (*n* = 7031 from six independent experiments) showed the expected GFP fluorescence at a level that could be determined visually. Among them, we could select colonies with the inserted target with 100% probability (*n* = 96) (Fig. 1D). Therefore, this TED-based expression system is an effective tool for both negative and positive colony screening.

### A TED-module-combined shuttle vector for magnetotactic bacteria

We applied the TED-based screening system to create a shuttle vector between *E. coli* and another prokaryote. A previous study reported a vector combined with pUC19 and a cryptic plasmid isolated from the magnetotactic bacterium *Magnetospirillum magneticum* (20). Based on this, we designed a vector contains the fragment necessary for its replication in magnetotactic bacteria and *E. coli*, a selection marker, and a fragment of the TED module inserted into the MCS (Fig. 2A and Supplementary Fig. S1).

**Figure 2.**
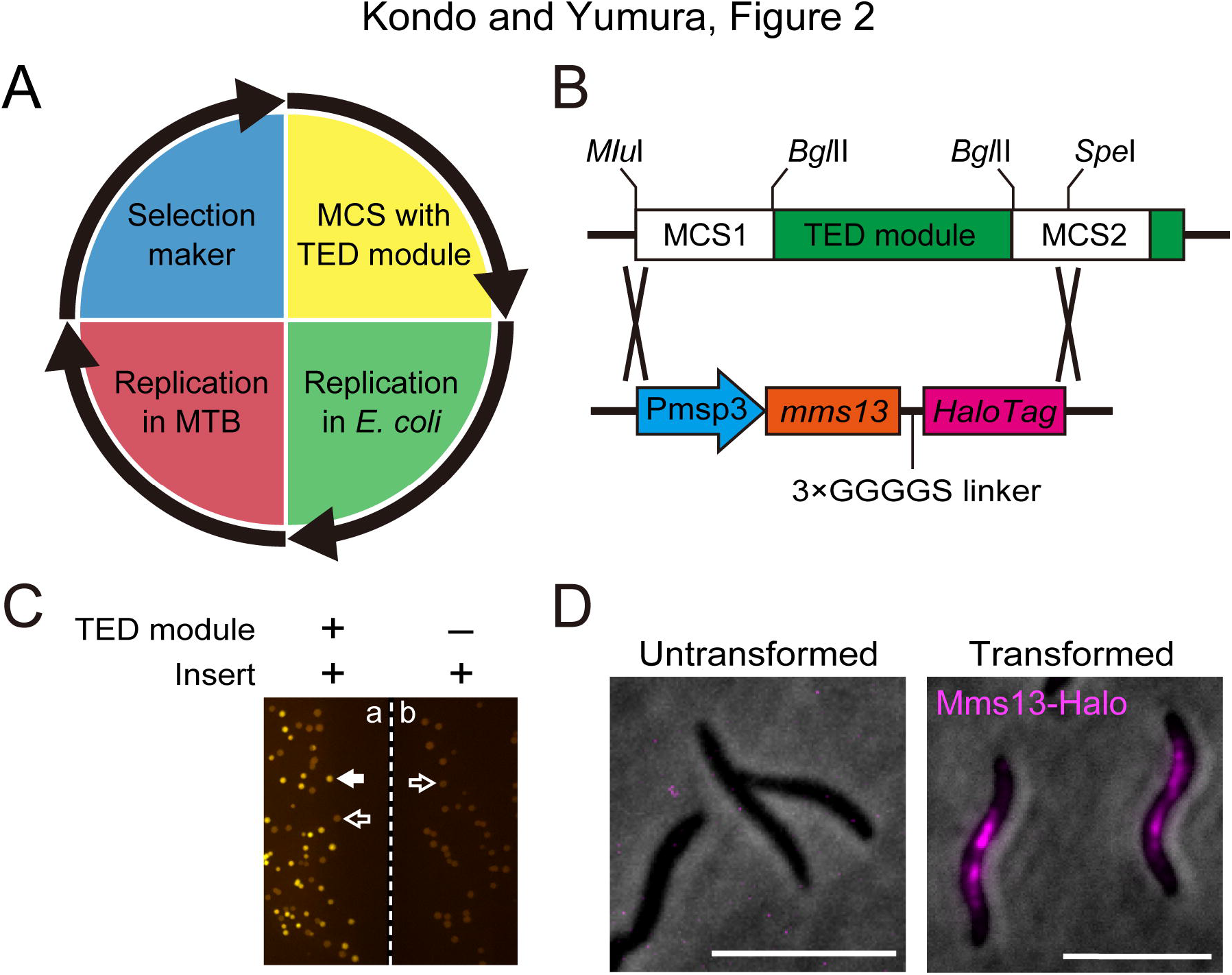
Negative screening of a shuttle vector for magnetotactic bacteria and visualization of their magnetosomes. (A) Schematic overview of the shuttle vector with four modules shown in different colors: yellow = MCS with the TED module; green = *E*. *coli* replication module; red = magnetotactic bacteria replication module; and blue = selection marker. (B) The sequence containing the *msp3* promoter (Pmsp3), *mms13*, and the HaloTag gene was inserted into the *Mlu*I-*Spe*I site. (C) Representative image of *E. coli* colonies cultured on an agar plate in the presence (a) or absence (b) of the TED module. Representative colonies with (white arrow) or without (open arrows) GFP expression are marked. (D) Image of magnetotactic bacteria MSR-1 stained with the HaloTag-TMR ligand. Bars, 5 µm.

The TED module was replaced with nucleotides encoding the magnetosome protein Mms13-fused HaloTag under control of the *msp3* promoter (P*msp3*) (22) to visualize magnetosomes (Fig. 2B). The *E. coli* colonies carrying plasmids with the desired substitution were selected by GFP-negative screening (Fig. 2C). Subsequently, the plasmid was transferred into a similar *Magnetospirillum* strain MSR-1, and the transformed cells were observed. The fluorescence of HaloTag ligands in MSR-1 cells was shown as aligned dot signals along the long axis of the cell at the midline (Fig. 2D), where the magnetosome chain is typically localized (28, 29). The *mms13* gene used here is derived from the AMB-1 strain and is known as *mamC* in the MSR-1 strain (80% amino acid sequence identity). Our observations revealed that Mms13 is still normally targeted to the magnetosomes, even in the MSR-1 strain.

To further test functionality and stability of our plasmid, we cultured cells in the presence or absence of antibiotics. The signals of HaloTag ligands that bind to HaloTag in cells were detected by image processing (Supplementary Fig. S2A). We found that our plasmid could function (Supplementary Fig. S2B) and be isolated (Supplementary Fig. S2C), even from cells cultured in an environment without selective pressure for more than 3 weeks. Considering that the doubling time of bacteria cultured in media is approximately 8 h, the plasmids were stably maintained in the strain. Therefore, we concluded that our shuttle vector functions properly for protein expression in magnetotactic bacteria.

### A TED-module-combined shuttle vector for *D. discoideum*

Next, we created a shuttle vector for eukaryotic *D. discoideum* in combination with the TED module. This vector consists of four components that are required for replication and selection in *Dictyostelium* or *E. coli* and expression of the target protein (Fig. 3A and Supplementary Fig. S3). The TED module was inserted into the MCS placed between the *Dictyostelium actin15* promoter and terminator to prepare a system capable of GFP-negative screening (Fig. 3B).

**Figure 3.**
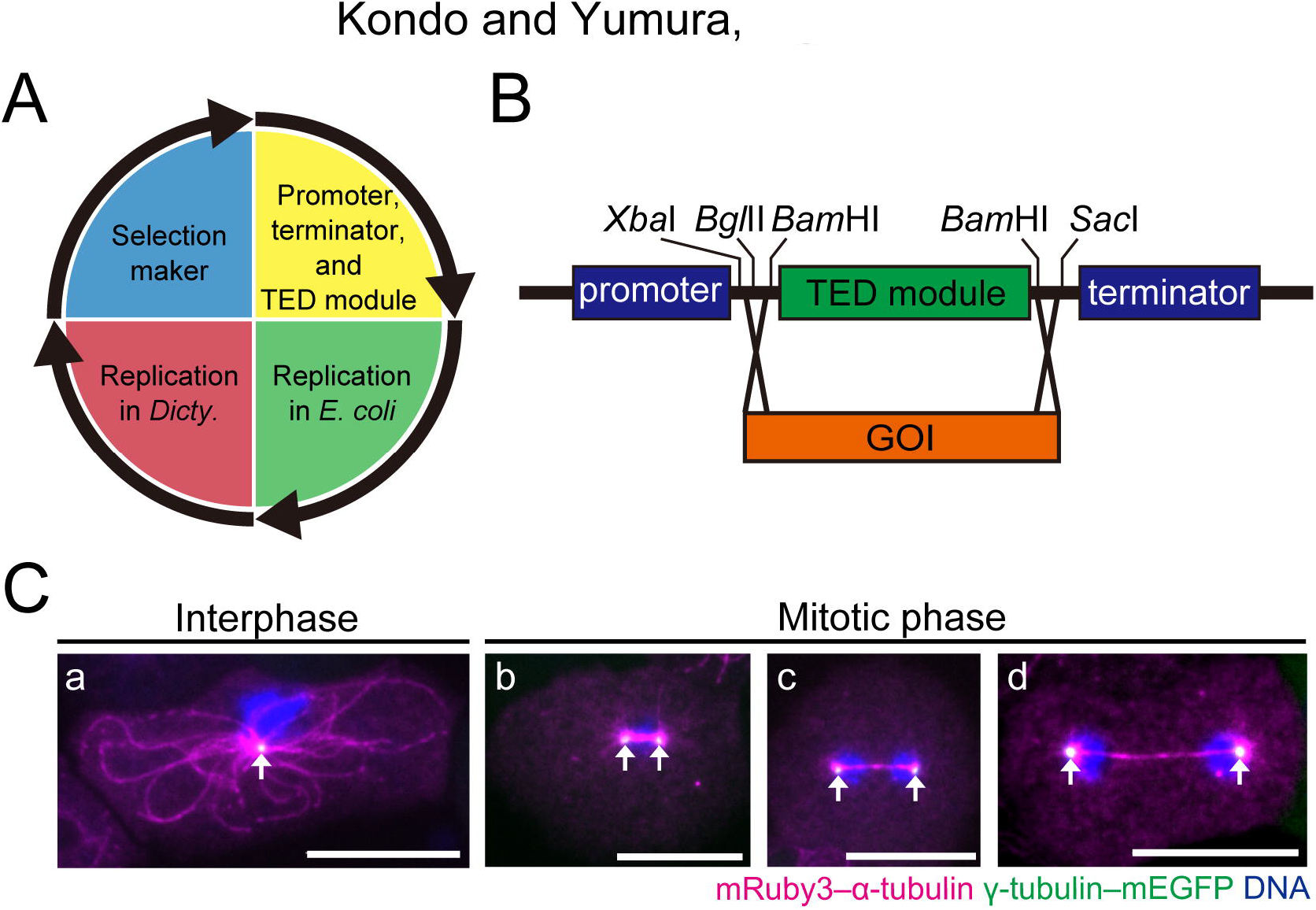
Negative screening of a shuttle vector for *Dictyostelium* and visualization of their cytoskeletal proteins. (A) Schematic overview of the extrachromosomal vector with the four modules shown in different colors: yellow = *actin15* promoter and terminator spacing by the MCS with the TED module; green = *E. coli* replication module; red = *Dictyostelium* replication module (a 2033 bp *Hind*III fragment of Ddp1); and blue = selection marker for *E. coli* and *Dictyostelium*. (B) The sequence encoding the GOI was inserted into the *Xba*I-*Bam*HI site between the promoter and terminator. (C) Fluorescence images of fixed cells co-expressing mRuby3–α-tubulin and γ-tubulin–GFP. Arrows indicate the centrosome. (a) Interphase, (b) metaphase, (c) anaphase, and (d) telophase. Bars, 10 µm.

We created shuttle vectors to express well-known proteins. Alpha-tubulin and γ-tubulin are extensively characterized proteins that constitute microtubules and centrosomes, respectively, in various organisms, including *Dictyostelium* (30–35). Two vectors were prepared by replacing the TED module with these target genes and genes encoding fluorescent proteins connecting a flexible (Gly-Gly-Gly-Gly-Ser) linker sequence (36). After bacterial colony screening and transformation into *Dictyostelium* cells, localization of the expressed protein was observed. As expected, mRuby3–α-tubulin showed a network structure, and γ-tubulin–GFP localized at the center of the network during interphase and mitosis, suggesting that the created vectors were functional (Fig. 3C).

Our vector was designed for simultaneous expression of multiple proteins from a single plasmid, which can reduce the time to generate transformants and suppress variations in their expression levels (37). We introduced more than one desired GOI into the vector using In-fusion cloning, which has high precision for cloning (Supplementary Fig. S1). A generated vector contained multiple GOIs arranged in tandem, separated by a promoter or terminator (Supplementary Fig. S4A). The gene encoding mRuby3–histone H1 (GOI2) (38, 39) was additionally incorporated into the plasmid containing the first gene encoding γ-tubulin–GFP (GOI1) with the promoter and terminator. Similar to a cell with the single expression vector, the centrosome was similarly visualized (Supplementary Fig. S4B and C). In addition, the nucleus was successfully visualized. Thus, our vector can be used for multiple protein expression after only one transformation. Taken together, our vectors can be used for heterologous protein expression in *Dictyostelium*.

## Conclusion

Here, we present a GFP-based method for screening colonies with the desired insert by inserting the TED module into various vectors. This new green-white screening method will aid in the routine construction of many vector types used in biological science studies. Thus, using TED with a wide variety of vectors, it is possible to provide a user-friendly process, construct many vectors with less effort, and finally increase the efficiency of experiments.

## Supplementary information

Supporting data of this manuscript are provided as a separate PDF file. The PDF file includes Supplementary Figures S1–4, and captions of the figures.

## Acknowledgements

We thank Drs. Yoshihiro Fukumori and Azuma Taoka (Kanazawa University, Japan) for providing protocols for handling magnetotactic bacteria and Dr. Yuki Hara (Yamaguchi University, Japan) for use of his microscope. Codon optimization of the nucleotide sequence that encodes the HaloTag protein was performed with permission from Promega Corporation. TK was supported by the Japan Society for the Promotion of Science Research Fellowships for Young Scientists. This research was supported by the Japan Society for the Promotion of Science [KAKENHI Grant Number 16J08310 and 19K15809 to TK].

## Competing interest statement

The authors declare no competing financial interests.

