## Supplemental Figures for "An improved molecular tool for screening bacterial colonies using GFP expression enhanced by a *Dictyostelium* sequence"

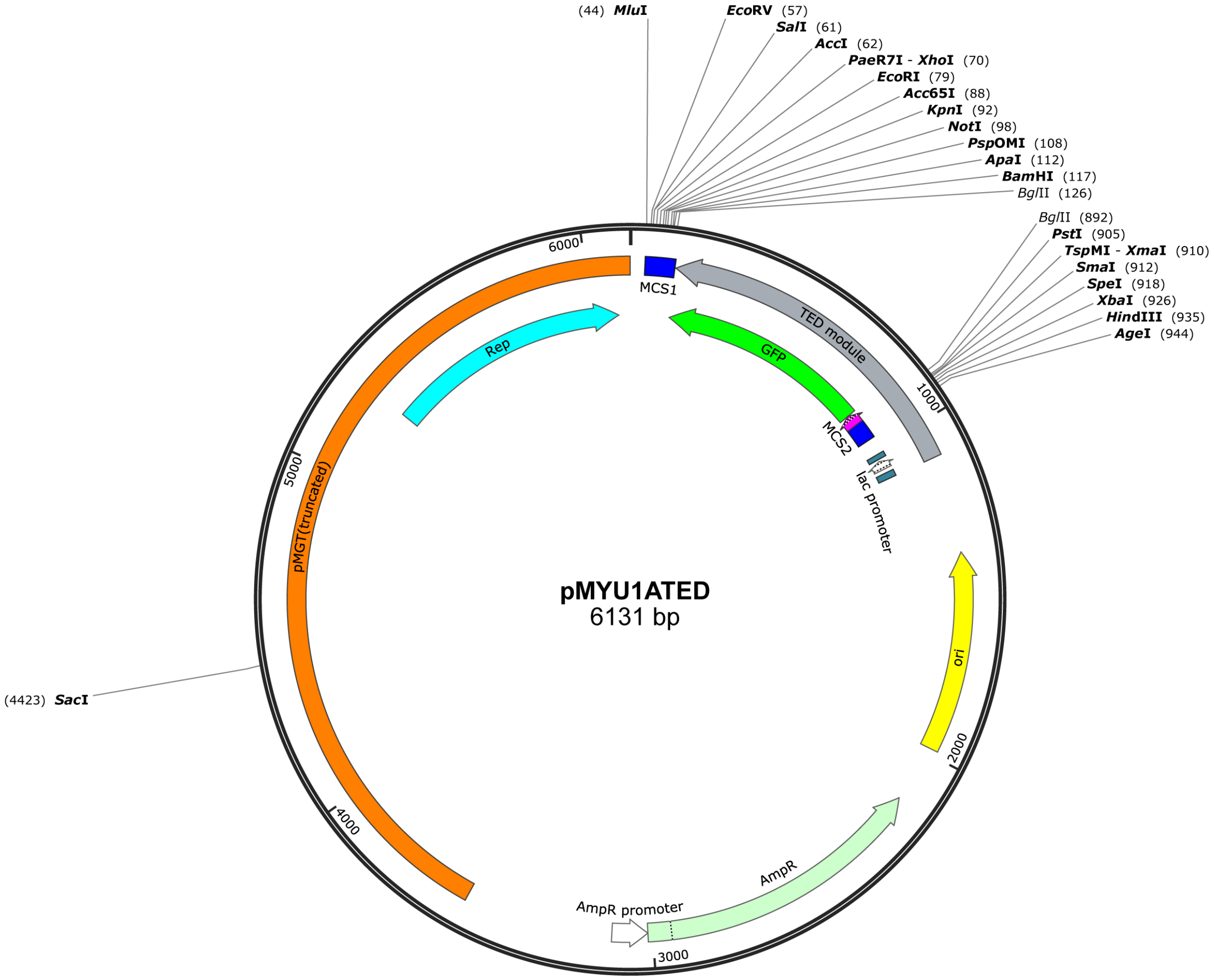


**Supplementary Figure S1. Plasmid map of pMYU1ATED.** pMGT (truncated) indicates a fragment of 2576 bp from position 18 to 2593 of pMGT (GenBank accession number NC_007706). The TED module is separated by MCS2 between *mlcR25* (magenta) and the *lac* operator (not labeled) to mimic the plasmid used in the original work (20).

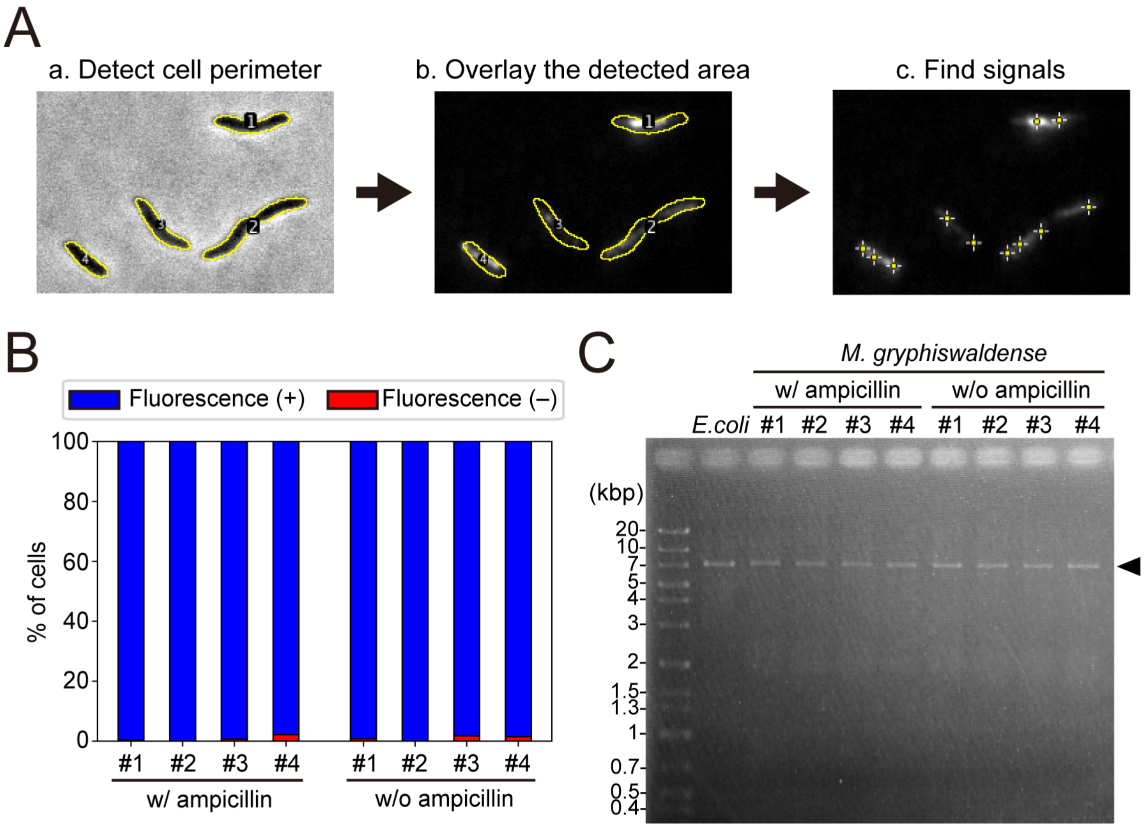


**Supplementary Figure S2. Quantification of magnetosome signals and plasmid stability in MSR-1.** (A) Example of image processing to detect signals from magnetotactic bacteria. Briefly, cell perimeters were detected from a phase-contrast image (a). The detected boundaries were overlaid on the fluorescent image (b). Finally, the signals were detected only inside the cell (c). (B) Quantitative analysis of cells cultured for 25 days in the presence or absence of ampicillin. Cells were passaged every 2–3 days. Each number (#1–4) represents a cell line. Fluorescence (+) indicates cells with HaloTag ligand signals. $n \geq 146$. (C) Image of an 1% agarose gel after electrophoresis of purified plasmids. *E. coli* indicates the lane with plasmids purified from *E. coli* cells. Each number (#1–4) represents a cell line of magnetotactic bacteria, which are the same ones shown in (B). All plasmids were linearized by a restriction enzyme reaction using *Mlu*I. The arrow indicates the predicted length of DNA bands. The gel edge was trimmed by image processing but no band enhancement was performed.

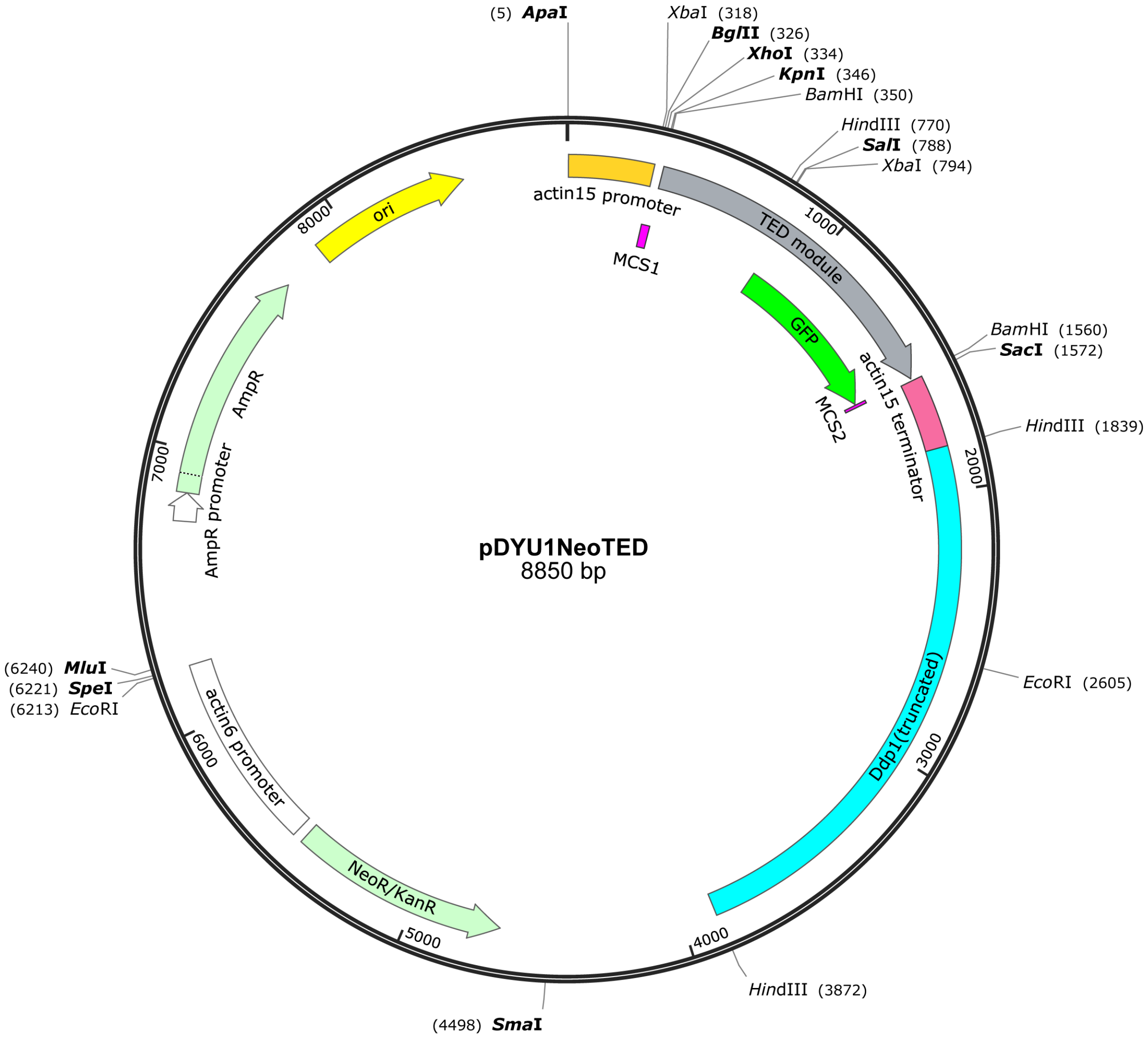


**Supplementary Figure S3. Plasmid map of pDYU1NeoTED**. Ddp1 (truncated) indicates a 2033 bp *Hind*III fragment of the pBIG vector (GenBank accession number AF270470). The TED module (1214 bp) contains 287 bp of the sequence upstream of the CAP binding site of pUC19. NeoR/KanR was replaced with *bsd* to create blasticidin S-resistant cell lines.

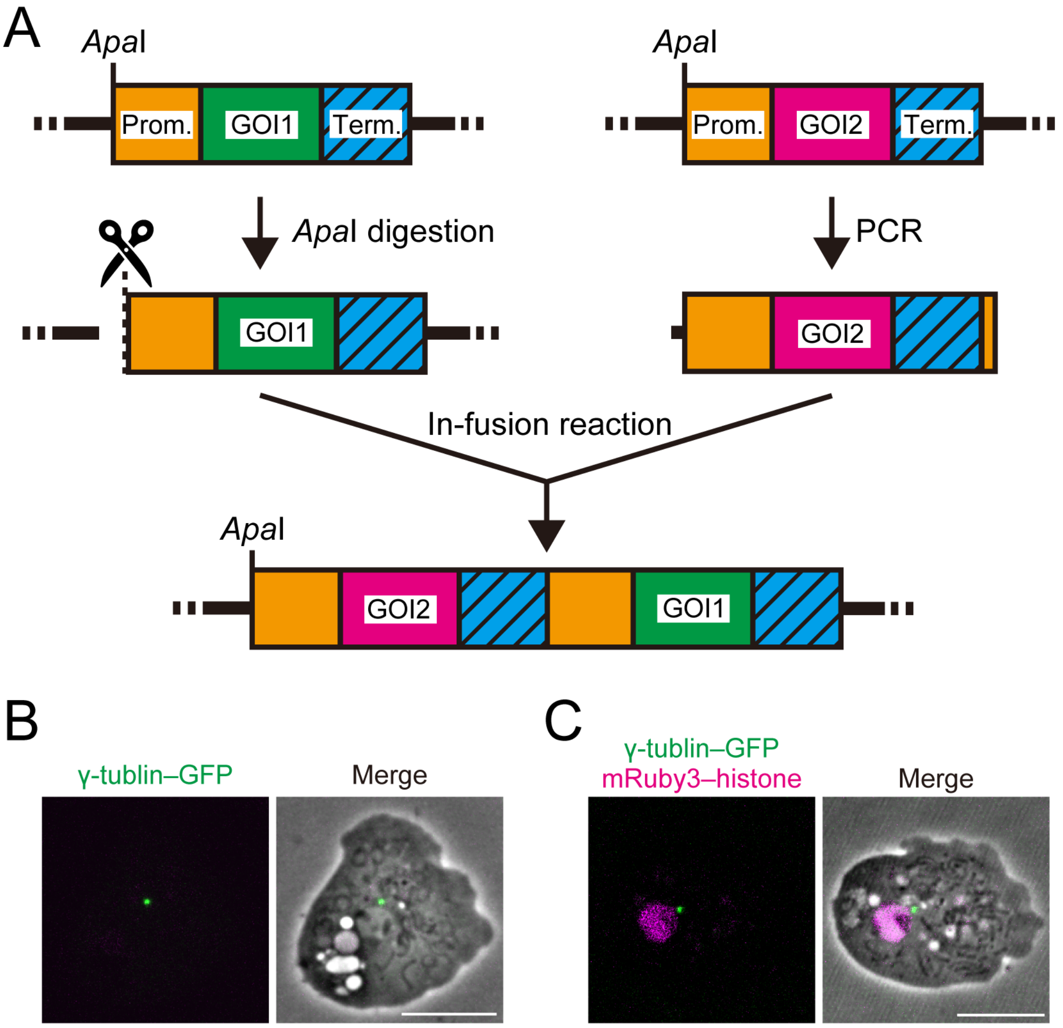


**Supplementary Figure S4. Simultaneous multiple protein expression from a single vector for *Dictyostelium.*** (A) Scheme of tandem GOI alignment using seamless cloning (In-fusion cloning). After linearization of the vector containing GOI1 with *Apa*I (whose recognition site is not present in 99.5% of the *Dictyostelium* genes), a PCR product containing GOI2 with a promoter (orange) and terminator (cyan, shaded) was inserted. (B) A γ-tubulin–GFP expressing cell. G418 was used for selection. (C) A γ-tubulin–GFP and mRuby3–histone H1 expressing cell. Only G418 was used for selection. The right side of each image superimposed to the phase-contrast image. Bars, 10 µm.
